# A Next Generation Clustering Tool Enables Identification of Functional Cancer Subtypes with Associated Biological Phenotypes

**DOI:** 10.1101/175307

**Authors:** Gift Nyamundanda, Katherine Eason, Anguraj Sadanandam

**Affiliations:** The Institute of Cancer Research, London, United Kingdom

## Abstract

One of the major challenges faced in defining clinically applicable and homogeneous molecular tumor subtypes is assigning biological and/or clinical interpretations to etiological (intrinsic) subtypes. The conventional approach involves at least three steps: Firstly, identify subtypes using unsupervised clustering of patient tumours with molecular (etiological) profiles; secondly associate the subtypes with clinical or phenotypic information (covariates) to infer some biological meaning to the redefined subtypes; and thirdly, identify clinically relevant biomarkers associated with the subtypes. Here, we report the implementation of a tool, phenotype mapping (*phenMap*), which combines these three steps to define *functional subtypes* with associated phenotypic information and molecular signatures. *phenMap* models meta (unobserved) variables as a function of covariates to expose any underlying clustering structure within the data and discover associations between subtypes and phenotypes. We demonstrate how this tool can more avidly identify functional subtypes that are an improvement over already existing etiological subtypes by analysing published breast cancer gene expression data.

## Background

A majority of the cancer types are heterogeneous representing a collection of diseases with different prognosis, responses to treatment, cellular-of-origin, molecular, metabolic and micro-environmental changes^1,2,3,4,5,6,7^. To define and implement clinically applicable tumor subtypes with associated biomarkers, it becomes important to integrate molecular profiles (genomics) with phenotypic information (including clinical outcome data such as tumor grade, stage and drug response; phenomics). This integrative stratification of tumors using genomics and phenomics is based on the principle that any phenotypes such as tumour grade and cancer cell survival are built upon interactions between multiple macromolecules such as DNA, RNA, proteins and metabolites. Studying these interactions thereby linking genomes to phenotypes forms a systematic way to understand the functional characteristics of patient tumors. Hence, there is a genuine need for robust statistical tools to integrate these molecular profiles with phenomics data to capture all information for effective implementation of subtypes in the clinic. In this study we are interested in combining a single molecular profile (gene expression) with associated multiple phenotypic data to identify subtypes and their associated phenotypes and biomarkers.

Recent initiatives such as The Cancer Genome Atlas (TCGA)^8^ and International Cancer Genome Consortium (ICGC)^9^ have lead to the generation of hitherto unprecedented levels of data on tumor genomics, metabolomics and phenomics. The conventional approach of tumor stratification involves unsupervised clustering approaches such as self-organizing maps (SOM)^10^, hierarchical clustering (HC)^11^, and non-negative matrix factorization (NMF)^3,6,12,13^. This unsupervised clustering approach uses only single or multiomics profiles (as in iCluster^14,15^) to identify subtypes that have distinct features reflective of different biology. Currently, associating the phenotypic data such as clinical information to the new subtypes is performed as a second step after clustering. Class prediction methods such as statistical analysis of microarrays (SAM) and predictive analysis of microarrays (PAM) are then used to identify differentially expressed features (genes, microRNA, etc), which stratify patients into different subtypes. This multi-stage approach can result in loss of statistical power to identify features driving the clustering or to discover association between subtypes and phenotypes. To address this issue we are proposing a more flexible approach to clustering through, phenotype mapping (*phenMap*), which combines these multiple steps to delineate “functional’’ subtypes (i.e. subtypes whose biological meaning and clinical implications are known).

## Results

### Application of *phenMap* in cancer

The *phenMap* framework can be applied to uncover clustering structure underlying the expression data, discover associated phenotypes and molecular features (i.e. genes, mRNAs, metabolites, etc.). The workflow of *phenMap* as a clustering framework, shown in Figure 1, requires profile data as an input, i.e., gene expression, along with associated phenotypes for a set of samples. Firstly, the framework identifies the optimal number of metavariables (MVs), *q*, driving the clustering of samples within the data using the Bayesian information criterion (BIC). Secondly, a fully Bayesian approach is taken to fit the model with the optimal number of MVs set at *q*. This results in i) clustering of samples in the *q*-dimensional MV space, ii) a panel of features driving this clustering and iii) associated covariates (phenotypes). Finally, model based clustering (MBC)^16^ is used to assign samples into their respective subtypes. Overall, *phenMap* model, unlike standard clustering approaches, identifies subtypes that are associated with sparse molecular signatures and biological phenotypes (covariates), hence, termed as “functional subtypes”. A complete description of the approach is available in methods and supplementary information sections.

**Figure 1:**
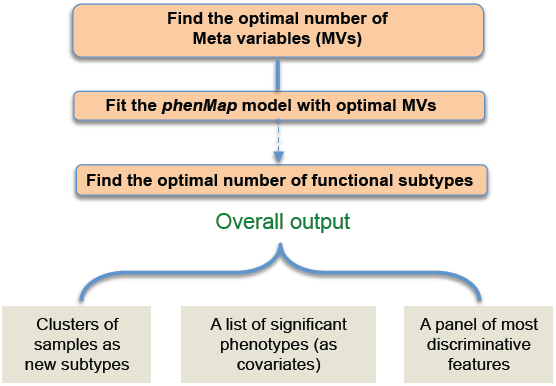
Modeling high dimensional data using *phenMap* model. A flowchart highlighting several steps involved in the *phenMap* framework when clustering samples.

### Functional gene expression subtypes of breast cancer

Here, we demonstrate the potential utility of *phenMap* in identifying functional subtypes in cancer using published gene expression data of 36 breast cancer cell lines profiled for different cancer drugs^17^. The top most variable 996 genes (standard deviation (SD)>1.2) were selected from 36 BrCa cell lines with matched drug inhibitory concentration (-log_10_ [IC_50_]) values for four therapeutic compounds; etoposide, fascaplysin, bortezomib and geldanamycin that were used as covariates (phenotypes).

When applied to this data, the model identified two MVs (Figure 2A) that clustered breast cancer cell lines into five “global (G) subtypes” (Figure 2B-C). Simultaneously, the model identified significant drugs (p<0.05) – fascaplysin (cyclin-dependent kinase; CDK inhibitor) and etoposide (chemotherapy) as well as important genes significantly (p<0.05) associated with the MVs, which can be further associated to subtypes using the magnitude and direction of the standardized regression and loading coefficients, respectively (Figure 2D and E).

**Figure 2:**
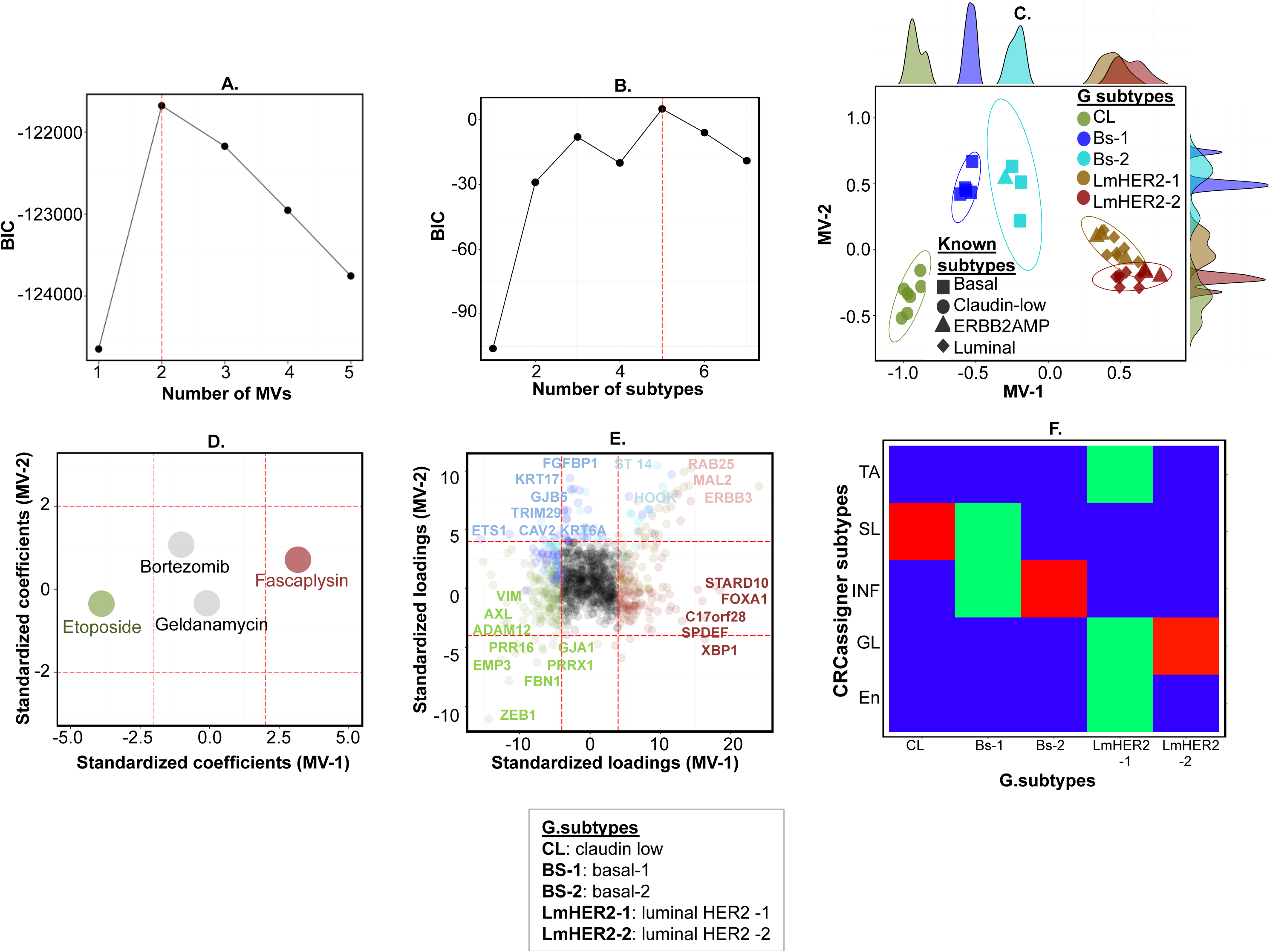
Applying *phenMap* to the gene expression data of breast cancer cell lines. **A**-**B.** Plots of BIC values to identify the optimal number of MVs and subtypes, respectively, for this dataset. The red dashed lines identify the optimal number of MVs and subtypes to be two and five, respectively. **C.** A plot showing subtypes in the MV space spanned by the two MVs. **D**-**E.** Plots showing D) the standardized regression coefficients (the red dashed lines represent 5% significance level) and E) the standardized loadings coefficients for LV 1 and 2 (the red dashed lines represent 0.01% significance level). The red dots denote covariates and genes with positive effect on LV-1, i.e. sensitive drugs and up-regulated genes in luminal-HER2 1 and 2. The green dots denote covariates and genes with negative effect on LV-1, i.e. sensitive drugs and up-regulated genes in claudin-low. **F.** Heatmap of hypergeometric test of enrichment between the five functional subtypes and the CRCassigner subtypes^3^ (towards red colour identifies significant association). The Heatmap shows that the claudin-low global functional subtype is a stem-like CRC subtype whilst luminal-HER2-2 and basal-2 are enriched for goblet-like and inflammatory CRC subtypes, respectively.

Interestingly, these five subtypes were associated with the three known breast cancer cell line subtypes^17^ (p<0.05; Fisher Exact test, Figure 2C). Whilst one of the G subtypes is clearly claudin-low subtype, interestingly, luminal and ERBB2^AMP^ subtypes combined into two different G-subtypes (named as luminal-HER2-1 and -2). In contrast, basal breast cancer cell lines (except for one ERBB2^AMP^ sample) split into two G-subtypes. Figure 2F shows that one of the basal subtype is enriched for inflammatory colorectal cancer (CRC) subtype^3^ (samples were assigned into CRC subtype with maximum Pearson correlation); hence it was named as inflammatory G-subtype. The magnitude and direction of MVs in Figure 2C and 2E clearly show that the gene characteristics of basal and inflammatory subtypes are in between the other three – claudin-low, luminal-HER2-1 and luminal-HER2-2 - breast cancer subtypes.

Moreover, subtype-specific genes were further refined using prediction analysis of microarrays (PAM) analysis (dubbed as breast cancer G-Assigner “BrCa-G-Assigner” gene signature; Figures 2E). We also applied our probabilistic version of principal component analysis with covariates (PPCCA)^18^ and NMF^3,6,12,13^ to this dataset. Although PPCCA cannot be used for subtype discovery, we found that etoposide and fascaplysin were associated with the clustering in the first principal component (this finding is similar to what we have shown with *phenMap*). NMF identified three subtypes that correspond to already known breast cancer intrinsic subtypes (data not shown). Hence, *phenMap* identifies unprecedented functional G-subtypes of breast cancer cell lines associated with drug response that were not revealed using other clustering tools.

## Discussion

Currently personalized approaches for diagnosis and treatment of many diseases including cancer require stratification of patients into sub-groups based on high-throughput molecular and phenotypic data. The conventional clustering approaches are employed to identify subtypes in omics data and later, phenotypes are associated with the subtypes as a post-clustering step. In the past two decades, this approach to subtyping has led to identification of subtypes in multiple cancer types that are clinically not in practice. Statistical power to discover association between the new subtypes and phenotypes is reduced due to several steps involved in this conventional subtyping approach. There is need for a new generation of clustering methods that can simultaneously cluster and integrate both omics and phenotypic data to identify subtypes of clinical importance.

Here we introduce a new concept of “functional subtyping” that involves identification of subtypes associated with phenotypes (covariates) using phenMap. This model can be applied to any continuous high-throughput omics data with matched covariates information to identify: i) subtypes, ii) covariates associated with the subtypes and iii) features driving the clustering. The application of this model in cancer is highlighted using a published omics data types (transcriptomics) with samples from breast cancer cell lines^17^. We compared the identified functional subtypes to already known and published breast cancer subtypes that were identified using the same data by conventional clustering approaches. This approach of functional subtyping offers the prospect of identifying robust and clinically important subtypes.

This model can also be extremely useful in other molecular datasets such as proteomics data. However, the current scope of phenMap model does not accommodate non-Gaussian data as profile data. Research on different link functions to allow phenMap to model count and categorical data is underway.

## Methods

### Datasets and samples

We used our already published gene expression microarray data for 36 breast cancer cell lines and four drug response (IC_50_) data^17^.

